# Plastid EF-Tu Regulates Root Development through Both the ATM Pathway and GUN1

**DOI:** 10.1101/2020.02.03.932574

**Authors:** Pengcheng Li, Junjie Ma, Xueping Sun, Chuanzhi Zhao, Changle Ma, Xingjun Wang

## Abstract

Impaired plastid translation affects various aspects of plant development, but the molecular mechanism remains elusive. Here, we described that the reduced function of plastid translation elongation factor EF-Tu encoded by *RAB GTPASE HOMOLOG 8D* (*Rab8d*) elicits defects in root development, including the reduced meristem size, programmed cell death (PCD) in the stem cell niche (SCN), and quiescent center (QC) division. We found that the ATAXIA-TELANGIECTASIA-MUTATED (ATM)-SUPPRESSOR OF GAMMA RESPONSE 1 module mediated overexpression of *SIAMSE-RELATED 5* in the root meristem region is responsible for the reduced meristem size in the *rab8d* mutant through arresting the cell cycle. The QC activation in *rab8d* is dependent on *ETHYLENE RESPONSE FACTOR 115*, which expression is tightly associated with the PCD in SCN. We further found that Rab8d physically interacts with GENOME UNCOUPLED 1 (GUN1), and GUN1 is required for inducing PCD in the *rab8d* SCN. However, the loss of GUN1 function in *rab8d* severely impairs the root architecture, suggesting that the GUN1-mediated renewal of stem cells is essential for maintaining root growth. Our observations extend our knowledge on the roles of ATM and GUN1 in regulating root development through mediating plastid translation dependent signals.

**One-sentence summary:** The *rab8d*-dependent plastid signal mediated by ATM and GUN1 regulates the root meristem size and renewal of root stem cells, respectively.

## INTRODUCTION

Plant root development requires an incessant generation of different types of cells by the pluripotent stem cells. In the root apical meristem (RAM), the stem cell niche (SCN) is comprised of a group of (usually four) specialized organizer cells, which divide slowly and are called the quiescent center (QC) as a consequence, and their surrounding stem cells. The QC cells are generated from a small lens-shaped daughter cell of the hypophysis at the globular stage of embryogenesis (Scheres et al., 1994). Later, the mitotically inactive QC induces their adjacent cells to become the root stem cells (van den Berg et al., 1997).

The phytohormone auxin plays a fundamental role in the establishment of root SCN. At early stages of embryogenesis, the suspensor transports the maternal auxin to the apical cell through several transport facilitators (Moller and Weijers, 2009; Robert et al., 2013; Robert et al., 2018). These proteins including PINFORMEDs (PINs) are required for the auxin dynamic distribution in the embryo proper, which determines the architectures of embryo and future seedling (Friml et al., 2003). At the mid-globular stage, an auxin maximum is observed in the hypophysis and the uppermost suspensor cells (Friml et al., 2003). This maximum is key for the division pattern of hypophysis and the subsequent formation of root QC (Friml et al., 2002; Blilou et al., 2005). At postembryonic stages, the stable auxin maximum present in the root QC is considered to be a positional cue for the SCN (Sabatini et al., 1999).

In postembryonic roots, two parallel molecular pathways have been proven to be involved in the SCN maintenance, namely, the PLETHORA (PLT) and SCARECROW (SCR) pathways. The PLT proteins belong to the APETALA2 (AP2)-domain transcription factor family and have four members which function redundantly in root development (Aida et al., 2004; Galinha et al., 2007). Their distribution is gradient along the RAM region with a maximum in the SCN, which is important for determination of the RAM size (Galinha et al., 2007). The loss-of-function mutants of *PLTs* display an abnormal division pattern in the hypophysis of embryo and a severely impaired RAM formation at the seedling stage (Galinha et al., 2007). The SCR pathway involves the intercellular movement of SHORT ROOT (SHR) from stele to the *SCR* expression domain and the transcriptional regulation by the SHR-SCR module (Nakajima et al., 2001; Sabatini et al., 2003). *SCR* is specifically expressed in the QC, cortex/endodermis stem cells, and the entire endodermal cell layer. In these cells, SHR directly targets to the promoter region of *SCR* (Levesque et al., 2006), and the SCR protein in turn maintains the nuclear localization of SHR (Cui et al., 2007). SHR and SCR also forms a heterodimer with each other to regulate a common set of targets for the maintenance of the QC identity (Cui et al., 2007). In addition, SCR also interacts with the RETINOBLASTOMA-RELATED (RBR) protein, which plays a role in controlling the division rate of QC cells (Cruz-Ramirez et al., 2013). More importantly, a recent study has reported that the Class I teosinte-branched cycloidea PCNA (TCP) proteins combine the above two pathways through interacting with both PLTs and SCR (Shimotohno et al., 2018), which further reveals their synergistic function for the SCN specification.

Intriguingly, versus the longevity of QC, that of stem cells has been observed to be hypersensitive to some environmental stresses, such as chilling, UV, and ionizing radiation (Furukawa et al., 2010; Hong et al., 2017; Johnson et al., 2018). The SCN-specific programmed cell death (PCD) under these stress conditions is primarily mediated by the DNA damage response pathway which involves ATAXIA-TELANGIECTASIA-MUTATED (ATM) and SUPPRESSOR OF GAMMA RESPONSE 1 (SOG1) (Fulcher and Sablowski, 2009; Hong et al., 2017; Johnson et al., 2018). ATM is a phosphoinositide-3-kinase-related protein kinase and functions conservatively in response to DNA double-strand breaks (DSBs) among eukaryotic species (Garcia et al., 2003; Shiloh, 2006). SOG1 belongs to the plant-specific NAC [petunia No Apical Meristem (NAM), Arabidopsis Transcription Activation Factor (ATAF), Cup-Shaped Cotyledon (CUC)] transcription factor family and its function is analogous to the mammalian p53 (Yoshiyama et al., 2009; Yoshiyama et al., 2013). The DSBs-triggered signal induces the ATM-dependent hyperphosphorylation of SOG1 (Yoshiyama et al., 2013), which plays roles in cell cycle arrest, promoting DNA repair, and programmed death of stem cells under severe cases through governing the downstream gene transcription (Yoshiyama, 2016). This mechanism is also involved in maintaining genomic integrity during seed germination (Waterworth et al., 2016).

QC functions in the replenishment of stem cells through division when programmed death is occurred. Several pathways have been identified to play a role in controlling the division rate of QC, including the WUSCHEL (WUS)-RELATED HOMEOBOX 5 (WOX5), RBR (mentioned above), and ETHYLENE RESPONSE FACTOR 115 (ERF115) pathways (Sarkar et al., 2007; Cruz-Ramirez et al., 2013; Heyman et al., 2013). As a member of the WUS family, WOX5 is highly enriched in QC and plays a key role in maintaining root columella stem cells (Pi et al., 2015). Its loss-of-function mutant displays ectopic QC divisions and lack of columella stem cells (Zhang et al., 2015). RBR is the plant homolog of the mammalian RB protein and exerts a traditional function in inhibition of cell cycle (Wildwater et al., 2005). Unlike the easily detected and stable expression of *WOX5* and *RBR*, the *ERF115* expression can be occasionally detected in QC under normal growth conditions (Heyman et al., 2013). Nevertheless, *ERF115* is highly induced by wound or PCD in the root SCN and its overexpression leads to extra divisions of QC through the PSK signaling, which is essential for renewal of the injured cells (Heyman et al., 2016).

RAB GTPASE HOMOLOG 8D (Rab8d, also called Rabe1b) is the nuclear-encoded plastid translation elongation factor EF-Tu in Arabidopsis. Previous studies have revealed the roles of Rab8d in heat stress response and regulation of leaf margin development (Li et al., 2018; Liu et al., 2019). In this study, we further found that Rab8d was required for root development. The *rab8d*-dependent plastid signal could be transduced by ATM, GENOME UNCOUPLED 1 (GUN1), and ERF115, which might play roles in controlling the RAM size, the longevity of stem cells, and the QC activity, respectively. Besides, our results also extended our knowledge on the role GUN1 in root development, which is known to be involved in the control of chloroplast biogenesis through mediating plastid-to-nucleus retrograde signaling and plastid proteostasis (Tadini et al., 2016; Marino et al., 2019; Wu et al., 2019; Zhao et al., 2019).

## RESULTS

### The Reduced function of Rab8d affects primary root development

According to the gene annotation from The Arabidopsis Information Resource (TAIR), the gene of *Rabe1b* is renamed *Rab8d* in this study. It has been reported that an allele of *Rab8d* with a T-DNA insertion in its single exon exhibits a lethal phenotype (Li et al., 2018; Liu et al., 2019), suggesting that this gene is essential for plant growth. To further investigate the function of *Rab8d*, we obtained three different T-DNA insertion lines for this gene. The insertion sites are indicated in Figure 1A. *rab8d-3* is another allele that contains a T-DNA insertion in the exon. Unsurprisingly, no homozygous plants of *rab8d-3* were isolated. Nevertheless, in mature-green siliques, we found that approximately 23.2% seeds were albino (Supplemental Figure 1). Most embryos in these albino seeds were arrested at the globular stage, and many of them had enlarged suspensors (a typical phenotype accompanied with arrested embryos, Supplemental Figure 1). In contrast, the embryos in normal seeds (green or brown) were all developed into the bent-cotyledon stage or beyond (Supplemental Figure 1). Because the plants developed from the normal seeds were *rab8d-3/+* or wild type, suggesting that the albino seeds might be homozygous *rab8d-3*. These observations further confirmed that *Rab8d* is vital for embryo development.

**Figure 1.**
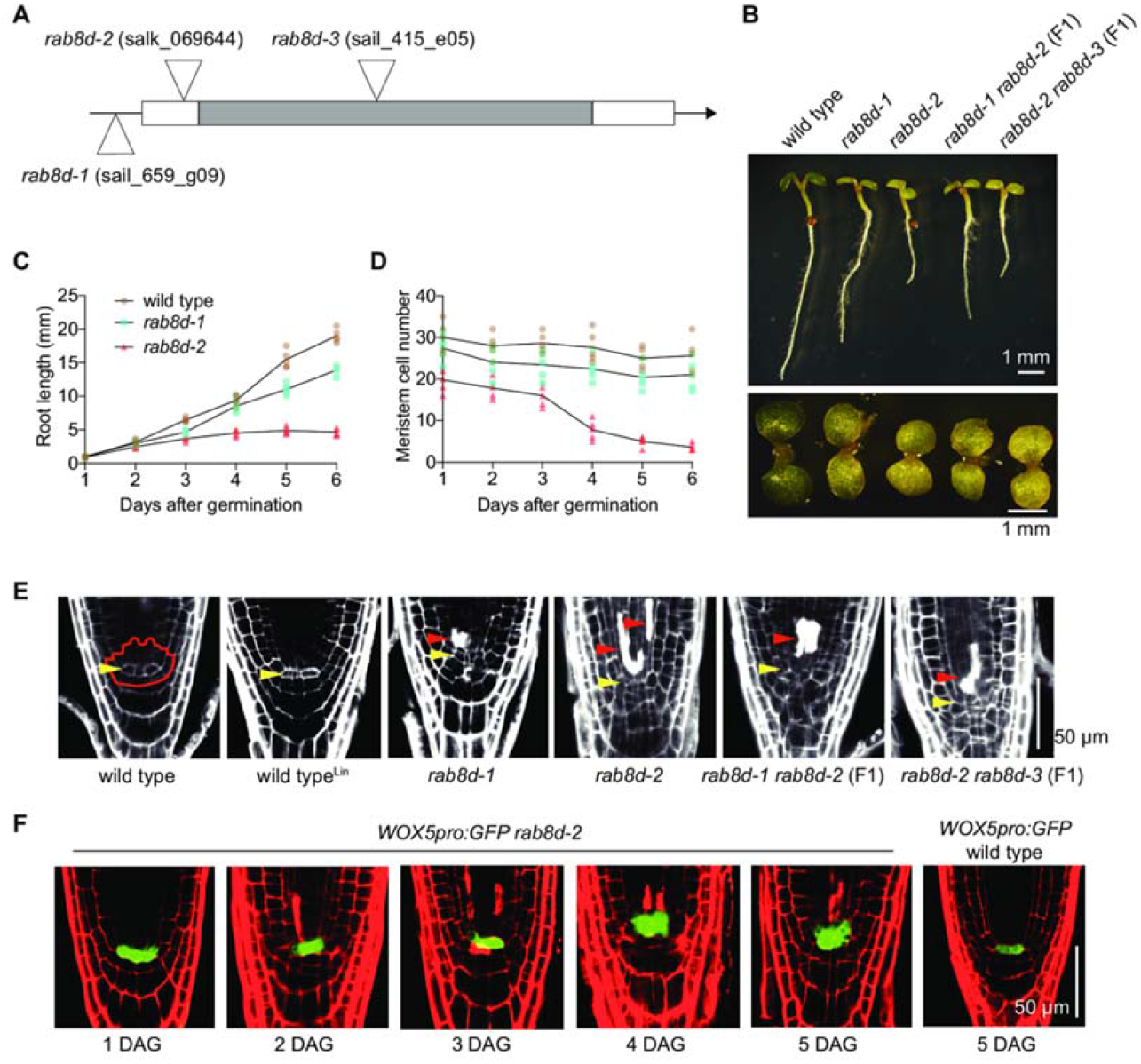
Root developmental phenotypes of the *rab8d* mutants. (A) Illustration of the *Rab8d* gene structures and positions of T-DNA insertions. The mutant names and corresponding accession numbers are indicated. (B) Phenotypes of 3 DAG seedlings of the indicated lines. (C) Kinematic analyses of primary root growth of the indicated lines from 1 to 6 DAG. (D) Dynamic changes of meristem cell number of the indicated lines from 1 to 6 DAG. (E) Phenotypes of SCN of the indicated lines. Wild type^Lin^ means a treatment with 220 mg/L lincomycin. Roots were stained with PI. The SCN is indicated in wild type with an irregular shape. Yellow arrowheads indicate the QC, and red ones indicate dead cells. (F) Expression patterns of *WOX5pro:GFP* in *rab8d-2* from 1 to 5 DAG. Bars are indicated.

The allele of *rab8d-1* (referred as *rabe1b-1* in a previous study) contains a T-DNA insertion in the gene promoter region (Figure 1A), which leads to an about three-quarter decrease in the gene transcriptional level (Li et al., 2018). *rab8d-2* contains a T-DNA insertion in the gene 5’ UTR (Figure 1A). Using quantitative PCR (qPCR), we further confirmed that *rab8d-1* retained about 30% of *Rab8d* mRNA levels in 5 DAG seedlings, whereas *rab8d-2* retained only about 8% of transcripts, indicating that both of them were knock-down alleles at least at the seedling stage (Supplemental Figure 2). The 3 DAG seedlings of both *rab8d-1* and *rab8d-2* displayed a pale-green phenotype under normal growth conditions (Figure 1B). Interestingly, the primary root of *rab8d-2* was significantly short than that of wild type (Figure 1B). The short-root phenotype was clearly mitigated in *rab8d-1* (Figure 1B), suggesting that the root length might be positively correlated with the *Rab8d* expression level. To further confirm this, we obtained two biallelic variants by crossing *rab8d-2* with *rab8d-1* and *rab8d-3*, respectively, *rab8d-1/rab8d-2* (F1) and *rab8d-2/rab8d-3* (F1). As displayed in Figure 1B, the root of *rab8d-1/rab8d-2* (F1) was shorter than *rab8d-1* but longer than *rab8d-2*, and the root of *rab8d-2/rab8d-3* (F1) appeared to be equal to *rab8d-2*. Notably, the *rab8d-2* root growth was arrested at 4 DAG (Figure 1C) with a rapid drawdown of the meristem cell number (Figure 1D). These results indicate that Rab8d is required for maintaining root growth. In addition, the retarded root growth phenotype of the *rab8d* mutants might not be associated with the photosynthesis in shoots because sucrose was applied in the growth medium as the carbon source.

More interestingly, when we visualize the outlines of root tip cells of 5 DAG seedlings with propidium iodide (PI, a fluorescent dye staining the walls of living cells), we found that some of the root initials of these *rab8d* mutants were filled with PI (Figure 1E), indicating the loss of membrane integrity and death of these cells. We also noticed that the cell death was occurred more frequently in stele initials and their daughter cells than in columella cells. Furthermore, the QC cells of *rab8d* were indistinguishable morphologically (Figure 1E), suggesting that these cells became active and were divided. To further confirm this, we tested the expression of a QC-specific marker, *WOX5pro:GFP*. Resulting from that *rabe1b-2* contained a low level of *Rab8d* transcripts and was viable, we employed it as the primary material. At 1 DAG, although no stem cells were taken up by PI and the QC cells could be easily identified, the GFP signal triggered by the *WOX5* promoter appeared to be slightly diffused into the stele initials (Figure 1F). At 2~3 DAG, the cell death was seen in some of the root initials with a similar expansion of the GFP signal (Figure 1F). Markedly, since 4 DAG, the *WOX5* promoter activity was expanded about two cell layers and was surrounded by dead cells (Figure 1F), suggesting division of the QC cells. As a control, in the wild-type background, no expansion of the *WOX5* expression was observed (Figure 1F). These results suggest that the root stem cells, especially stele initials, are hypersensitivity to the reduced function of Rab8d, and the PCD in SCN may further trigger the QC division for the stem cell replenishment.

In addition, resulting from that Rab8d is predicted to function in plastid translation, we wondered if the root growth defect of the *rab8d* mutants was associated with the impaired plastid translation. To test this, we treated the wild-type seedlings with a plastid translation inhibitor, lincomycin. The results showed that, though the root length was slightly reduced by lincomycin (Supplemental Figure 3A), no cell death was observed in SCN (Figure 1E) and QC was inactive indicated by the *WOX5* expression (Supplemental Figure 3B), suggesting that the Rab8d-dependent signal possessed a special role in fine regulating root development.

### Rab8d is highly accumulated in embryos and RAM

The subcellular localization of Arabidopsis Rab8d has been tested in leaf epidermal cells of tobacco (*N. benthamiana*) and Arabidopsis protoplast using transient expression by other labs, which results indicate that Rab8d is localized in chloroplast and formed distinct foci (Lichocka et al., 2018; Liu et al., 2019). We obtained the stable transgenic lines of *Rab8dpro:Rab8d-GFP* in the *rab8d-2* background. These transgenic plants were displayed a wild-type phenotype, suggesting that the fusion protein of Rab8d-GFP was fully functional *in vivo*. However, inconsistently, under normal growth conditions, we found that Rab8d-GFP was not formed foci but evenly distributed in chloroplast stroma of mesophyll cells (Supplemental Figure 4). A previous study has reported that the Rab8d protein is heat-sensitive because of its rapid aggregation at high temperatures (Li et al., 2018). We found that when these plants were treated with a heat-stress condition (37°C) for 4 h, agglomerated GFP signals were clearly observed in chloroplasts (Supplemental Figure 4). Therefore, our result may represent the true localization of Rab8d *in vivo*.

Since its vital roles in embryo development, the *Rab8d* expression pattern was analyzed in embryos of the *Rab8dpro:Rab8d-GFP* line. Indeed, Rab8d could be clearly detected throughout the embryo development process, even before the globular stage (Figure 2A). Note that no chlorophyll (Chl) fluorescence was detected in embryos at pre-globular and globular stages (Figure 2A), indicating that Rab8d was also present in proplastids. Together with the observation of albino seeds in the *rab8d-3+/-* siliques (Supplemental Figure 1), these results suggested that Rab8d might be required for plastid/chloroplast maturation during embryo development. The pattern of Rab8d protein abundance was further tested in roots, which result showed that its protein was highly accumulated in the RAM region, including in QC and stem cells (Figure 2B).

**Figure 2.**
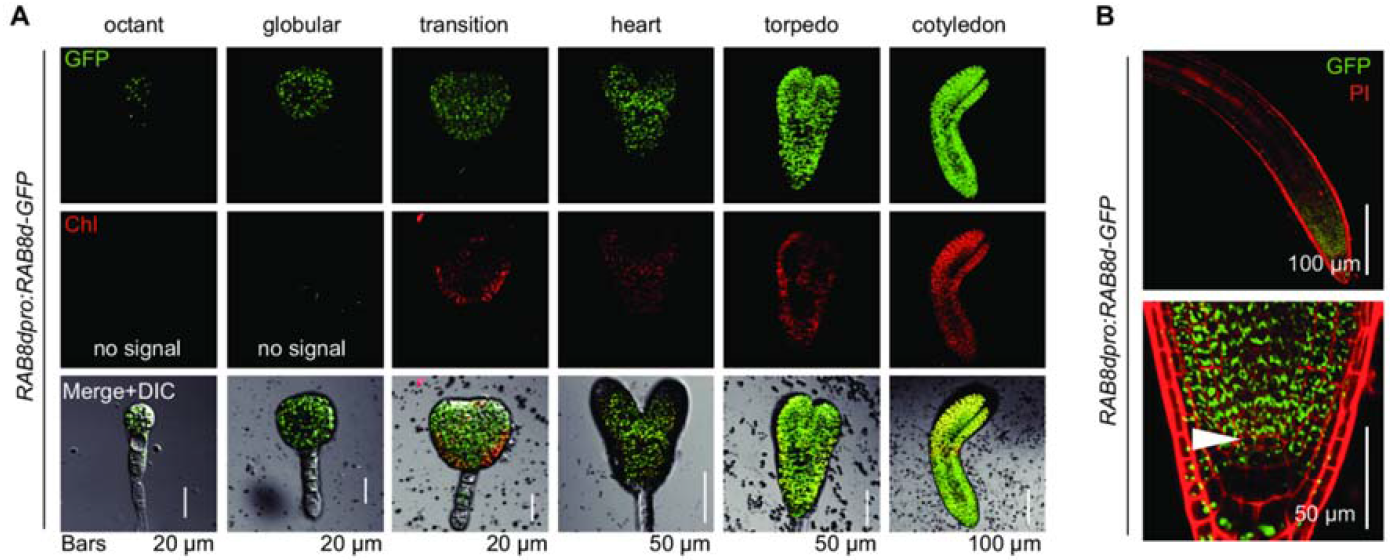
Expression patterns of *Rab8d* in embryos and roots. (A) Expression patterns of *Rab8d* in embryos from the octant stage to the cotyledonary stage. Chl, chlorophyll autofluorescence. (B) The accumulation of Rab8d protein in root tips of 3 DAG seedlings. White arrowheads indicate the QC. Bars are indicated.

### The loss of QC identity in *rab8d* correlates with the perturbed auxin maximum and PLT and SHR pathways

Auxin is required for the root meristem maintenance (Brumos et al., 2018). The retarded root growth resulting from the reduced level of Rab8d led us to estimate the auxin level in root tip. To address this issue, we employed an auxin-responsive marker *DR5rev:GFP* and tested its activity in root tips of wile type and *rab8d-2* using confocol. The results showed that, starting from germination, the auxin level (indicated by the GFP signal) in *rab8d-2* was obviously lower than that in wild type (Figure 3A). In QC, there is always an auxin maximum that is required for maintaining the cell identity (Sabatini et al., 1999). However, in *rab8d-2*, the GFP intensity peak in QC was undetectable (Figure 3B). This result is consistent with the observation that the QC cells lost their identity and are active in *rab8d-2* (Figure 1E, F). PINs-mediated polar auxin transport is critical for the establishment of auxin gradients (Friml et al., 2003; Blilou et al., 2005). Unsurprisingly, the accumulation of PIN1, PIN2, PIN3, and PIN7 proteins was also significantly reduced in the root of *rab8d-2* (Supplemental Figure 5). In addition to auxin, transcription factors of PLTs are also accumulated in gradient in root tip with maxima in QC and stem cells, which play key roles in the SCN patterning (Galinha et al., 2007). In *rab8d-2*, expression of *PLT1* and *PLT2* was severely downregulated, and the maximum of *PLT2* was lost, though that of *PLT1* could be detected weakly in QC (Figure 3C). The SHR-SCR module is also important for the SCN patterning, paralleling the PLT pathway (Nakajima et al., 2001; Sabatini et al., 2003). In the wild-type background, the SHR protein was clearly observed in the stele cells and nuclei of the QC and endodermal cells. In *rab8d-2*, SHR could be weakly observed in the stele cells, however, no (or very weak) signal was observed in QC or endodermal cells (Figure 3D). Consistently, the accumulation of its downstream factor, SCR, was also significantly decreased (Figure 3D). These results suggest that the loss of QC identity is tightly associated with the abnormal auxin distribution and the impaired PLT and SHR-SCR signaling pathways. The low levels of *PLTs* also positively correlate with the reduced RAM size in *rab8d-2*.

**Figure 3.**
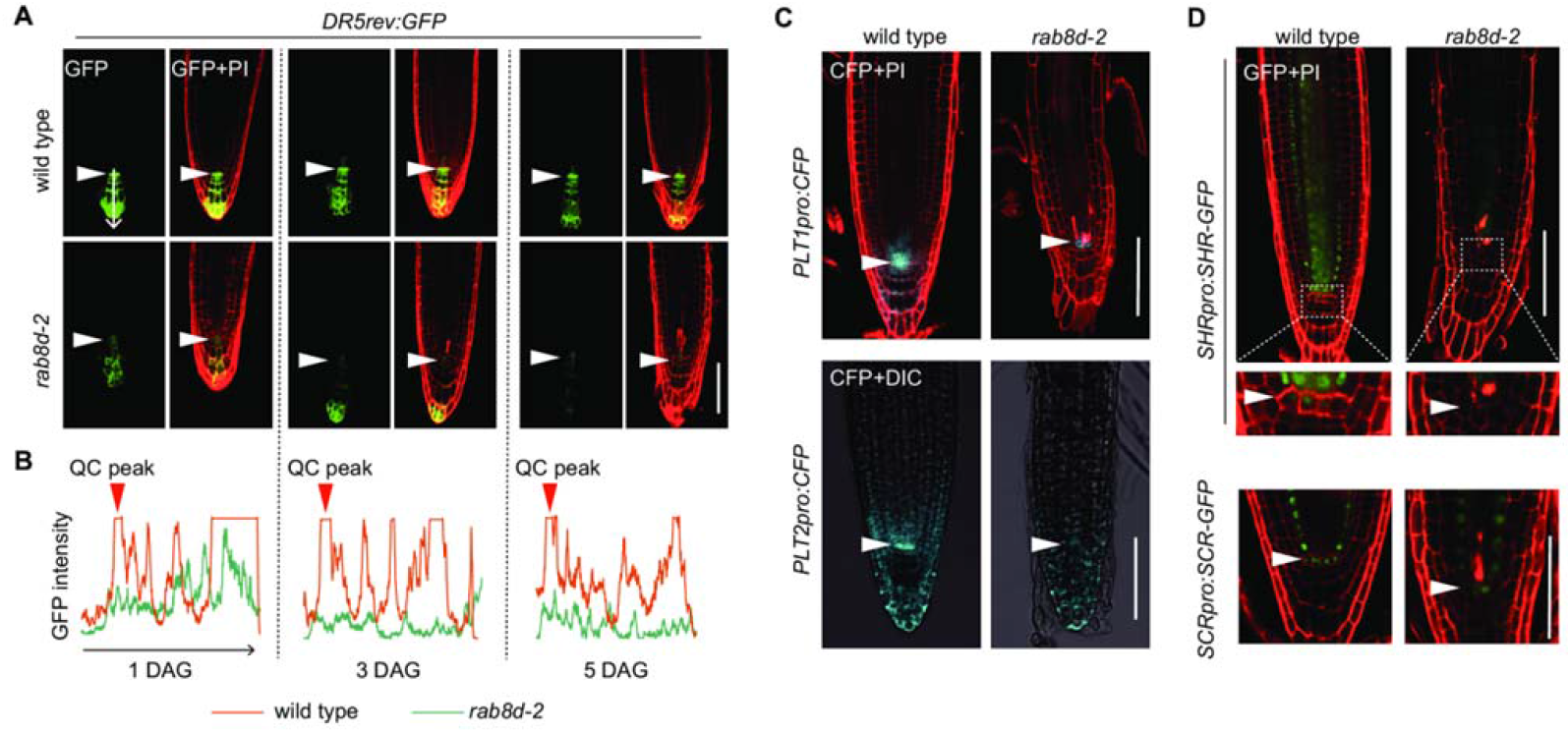
QC identity is lost in *rab8d-2*. (A) Expression of *DR5rev:GFP* in root tips of wild type and *rab8d-2* at 1, 3, and 5 DAG. (B) GFP signal intensity was measured with the software ImageJ according to the arrow indicated in (A). The QC peak indicates the auxin maximum in QC of wild type roots. (C) Expression of *PLT1pro:CFP* and *PLT2pro:CFP* in root tips of 5 DAG seedlings of wild type and *rab8d-2*. (D) Expression of *SHRpro:SHR-GFP* and *SCRpro:SCR-GFP* in root tips of 5 DAG seedlings of wild type and *rab8d-2*. White arrowheads indicate QC. Bars = 50 μm.

We further tested the root growth phenotype of *rab8d-2* grown under exogenous auxin conditions. The results showed that the exogenous auxin treatment (0.5, 1, and 2 nM IAA) had no clear effect on the *rab8d-2* root (Supplemental Figure 6A), suggesting that the retarded root growth of *rab8d-2* might not be resulted from the reduced auxin level *in vivo*. In contrast, the wild type roots were sensitive to the IAA treatment reflected by the reduced root length compared with control (Supplemental Figure 6A). Brassinosteroids also play roles in regulating root growth (Mussig et al., 2003). Note that a very low level of exogenous brassinolide (BL, 0.01 nM) application significantly promoted root growth of wild type (Supplemental Figure 6B). By contrast, the root growth of *rab8d-2* was further inhibited under the same condition (Supplemental Figure 6B), suggesting that *rab8d-2* was more sensitive to exogenous BL than wild type. Under higher concentrations of BL (0.05 and 0.1 nM), both wild type and *rab8d-2* displayed a decreased root length phenotype (Supplemental Figure 6B). Together, the retarded root growth of *rab8d-2* is likely independent of auxin or BR signaling.

### Root transcriptome comparison between wild type and *rab8d-2*

To further figure out the molecular mechanism of the regulation of root growth by Rab8d, we performed transcriptome profiling using RNA-sequencing (RNA-seq) to find out changes in global gene expression between wild type and *rab8d-2*. Total RNA was extracted from roots of 5 DAG seedlings, and a total of 24233 genes were detected (Supplemental Dataset 1). Consistent with the observation of increased *WOX5* promoter activity in *rab8d-2*, its transcript abundance was elevated about 3.5 folds (Supplemental Figure 7). The mRNA levels of *SHR, SCR, PLT1, PLT2* and *PIN2* were all decreased to some extent (Supplemental Figure 7). However, the transcript abundance of *PIN1* and *PIN7* was not clearly affected, and that of *PIN3* appeared to be slightly increased (Supplemental Figure 7), suggesting a potential posttranscriptional response of these genes to the defect of Rab8d. Compared to wild type, *rab8d-2* possessed 2587 up- and 2149 downregulated genes based on the fold change > 2 and adjusted *P* value < 0.001 (Supplemental Dataset 2). Through analyzing the Gene Ontology (GO) enrichment in terms of biological processes, we found that approximately 72 upregulated genes were associated with photosynthesis (Supplemental Dataset 3), suggesting that this process was sensitive to the reduced function of Rab8d, even in the non-photosynthetic tissue, the root. More interestingly, approximately a quarter of upregulated genes were associated with response to stimulus, including abiotic and biotic stresses and hormones (Supplemental Dataset 3). The result suggests that the reduced function of Rab8d mimics these stress conditions and affects homeostasis of their responsive gene transcripts. By contrast, the downregulated genes were primarily associated with signaling pathways including the PCD, protein modification, cell growth and cycle regulation, and root development (Supplemental Dataset 4).

### The reduced root meristem size in *rab8d* is partially rescued by perturbing the ATM pathway

Among the mis-regulated genes in *rab8d-2*, the dramatically increased transcript abundance of the gene encoding one of the cyclin-dependent kinase inhibitors, *SIAMSE*-*RELATED5* (*SMR5*, Figure 4A), attracted our attention because of its role in cell cycle inhibition (Yi et al., 2014). To further confirm this result, we compared the *SMR5pro:GUS* activity between wild type and *rab8d-2*. In the wild-type background, *SMR5pro:GUS* was predominantly expressed in the columella cells with a very low level in the meristem region (Figure 4B). However, in the *rab8d-2* background, the expression of *SMR5pro:GUS* was markedly observed in the meristem region (Figure 4B). We thus assumed that the cell cycle might be affected in the root meristem of *rab8d-2*. To test this hypothesis, we first compared the density of cells entered the S phase [detected with the thymidine analogue 5-ethynyl-2’-deoxyuridine (EdU)] between wild type and *rab8d-2* (see details in Methods). The results showed that the density of cells entered the S phase was reduced by roughly half in *rab8d-2* (Supplemental Figure 8A, B). Then we employed the *CycB1;1pro:GUS* reporter to indicate cells at the G2/M transition phase. Using GUS staining, we found that the density of GUS-positive cells was significantly increased in *rab8d-2* (Supplemental Figure 8C, D). On account of the affected S-phase entry, the increased G2/M cell density in *rab8d-2* suggested the lack of M-phase entry because of degradation of the fused protein of GUS with a mitotic destruction box at M phase. These observations suggest that Rab8d is important for the global cell cycle maintenance, and the reduced meristem size of *rab8d-2* may be resulted from the affected cell cycle. To further investigate whether the reduced meristem size of *rab8d-2* was associated with the induction of *SMR5*, we eradicated the SMR5 function in *rab8d-2* by crossing it with a knockout allele of *smr5*. Our observations indicated that the loss of SMR5 function indeed rescued, but not fully, the root growth phenotypes of *rab8d-2*, including the root length and the number of meristem cells (Figure 4C-E).

**Figure 4.**
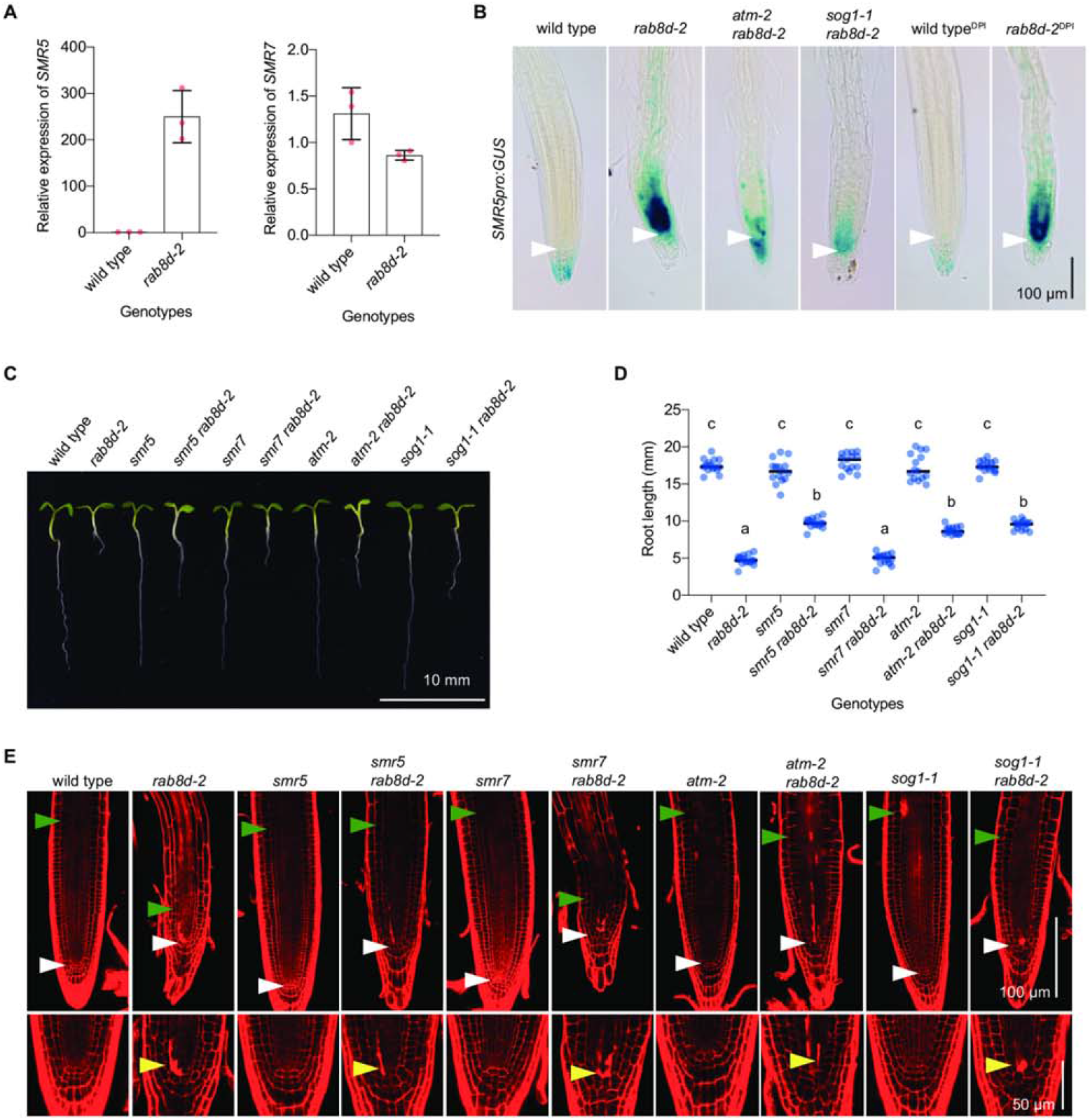
*smr5* partially rescues the meristem phenotype of *rab8d-2*. (A) Comparison of expression level of *SMR5* and *SMR7* between wild type (set to 1) and *rab8d-2*. The data was based on the FPKM values tested by RNA-seq. (B) The expression of *SMR5pro:GUS* in root tip of 5 DAG seedlings of wild type, *rab8d-2, atm-2 rab8d-2, sog1-1 rab8d-2*, and DPI-treated wild type (wild type^DPI^) and *rab8d-2* (rab8d-2^DPI^). White arrowheads indicate QC. (C) Phenotypes of 5 DAG seedlings of the indicated genotypes. (D) Comparison of root length among the indicated genotypes. Each spot represents an individual value, and black dashes indicate mean values. Significance analysis was performed using one-way ANOVA, and different lowercases indicate *P* < 0.0001. (E) Root tip phenotypes of 5 DAG seedlings of the indicated genotypes. White arrowheads indicate QC, green ones indicate the basal cell of the elongation region, and yellow ones indicate dead stem cells. The distance between white and green arrowheads indicate the meristem size.

We also noted that the expression pattern of *SMR5* in the *rab8d-2* root is very similar as observed in the root treated with the DNA replication inhibitory drug hydroxyurea (Yi et al., 2014). The induction of *SMR5* on the genotoxic stress conditions is dependent on ATM and SOG1 (Yi et al., 2014). As a consequence, we wondered whether the induction of *SMR5* in the *rab8d-2* root was also mediated by the ATM pathway. To address this issue, we introduced the *SMR5pro:GUS* reporter into the double mutant backgrounds of *atm-2 rab8d-2* and *sog1-1 rab8d-2*. Using GUS staining, we found that the loss of ATM or SOG1 function dramatically reduced the activity of *SMR5pro:GUS* in the RAM of *rab8d-2* (Figure 4B). The observation suggested that the ATM-SOG1 pathway could perceive the *rab8d*-dependent signal. In addition, the *SMR5* expression also can be highly induced by ROS, which is mediated by the ATM pathway as well (Yi et al., 2014). We further wondered whether the *SMR5* induction in response to the reduced function of *Rab8d* was associated with ROS. Using 3,3-diaminobenzidine (DAB) staining, we observed no significant change in the H2O2 level in *rab8d-2* relative to that in wild type (Supplemental Figure 9A). Yet, using nitroblue tetrazolium (NBT) staining, we found that the superoxide accumulation was higher in the root of *rab8d-2* than in that of wild type, especially in the stele of elongation zone (Supplemental Figure 9B). To investigate the role of superoxide in the regulation of *SMR5* in *rab8d-2*, we treated seedlings with an NADPH oxidase inhibitor, diphenylene iodonium (DPI). However, in *rab8d-2*, no obvious difference in the *SMR5pro:GUS* activity was observed between seedlings grown under control and DPI (Figure 4B), which indicated that the induction of *SMR5* expression was not associated with the ROS signaling. Consistently, DPI had no clear effect on the root growth phenotype of the *rab8d* mutants, whereas it slightly reduced the root length of wild type (Supplemental Figure 9C).

As *smr5* partially rescued the root phenotypes of *rab8d-2*, we further examined the effect of loss of ATM or SOG1 function on them. Compared with *rab8d-2*, both the double mutants of *atm-2 rab8d-2* and *sog1-1 rab8d-2* possessed significantly longer roots and more root meristem cells (Figure 4C-E), although these rescues were partial likewise. More importantly, the ATM-SOG1 pathway mediates the DNA damage induced PCD in SCN (Fulcher and Sablowski, 2009; Yoshiyama et al., 2013). However, the loss of ATM, SOG1 or SMR5 had no effect on the PCD phenotype in SCN of *rab8d-2* (Figure 4E).

In addition, consistent with *SMR5, SMR7* is also in response to the ATM and SOG1 mediated ROS signaling and plays a role in regulating cell cycle (Yi et al., 2014). However, based on the RNA-seq results, the abundance of *SMR7* transcripts was slightly reduced in the *rab8d-2* root (Figure 4A). Therefore, it is not surprised that the loss of SMR7 function had no effect on root development of *rab8d-2* (Figure 4C-E).

Together, the above results indicate that the ATM-SOG1 module perceives the *rab8d*-dependent signal and plays a role in regulation of RAM size in *rab8d-2* through inducing the *SMR5* expression.

### The loss of QC identity in *rab8d-2* is dependent on ERF115

The strong reaction of *ERF115* to the reduced expression of Rab8d (Figure 5A) also attracted our attention because of its roles in regulating the QC activity and stem cell renewal (Heyman et al., 2013; Heyman et al., 2016). Through detecting the GFP signals triggered by *ERF115pro:NLS-GFP* in the *rab8d-2* background, we found that the *ERF115* expression was highly induced in the meristematic stele cells surrounding the dead ones from 3 DAG, whereas no (or extremely weak) signals were observed in the wild-type background (Figure 5B). The hyperactive of *ERF115* expression might be associated with the death of stele initials because of its wounding-responsive characteristic (Heyman et al., 2016). Nevertheless, at 1.5 DAG, no cell death was observed but the *ERF115* expression could be clearly detected in SCN of *rab8d-2* (Figure 5B), suggesting onset of programmed death of some of these cells.

**Figure 5.**
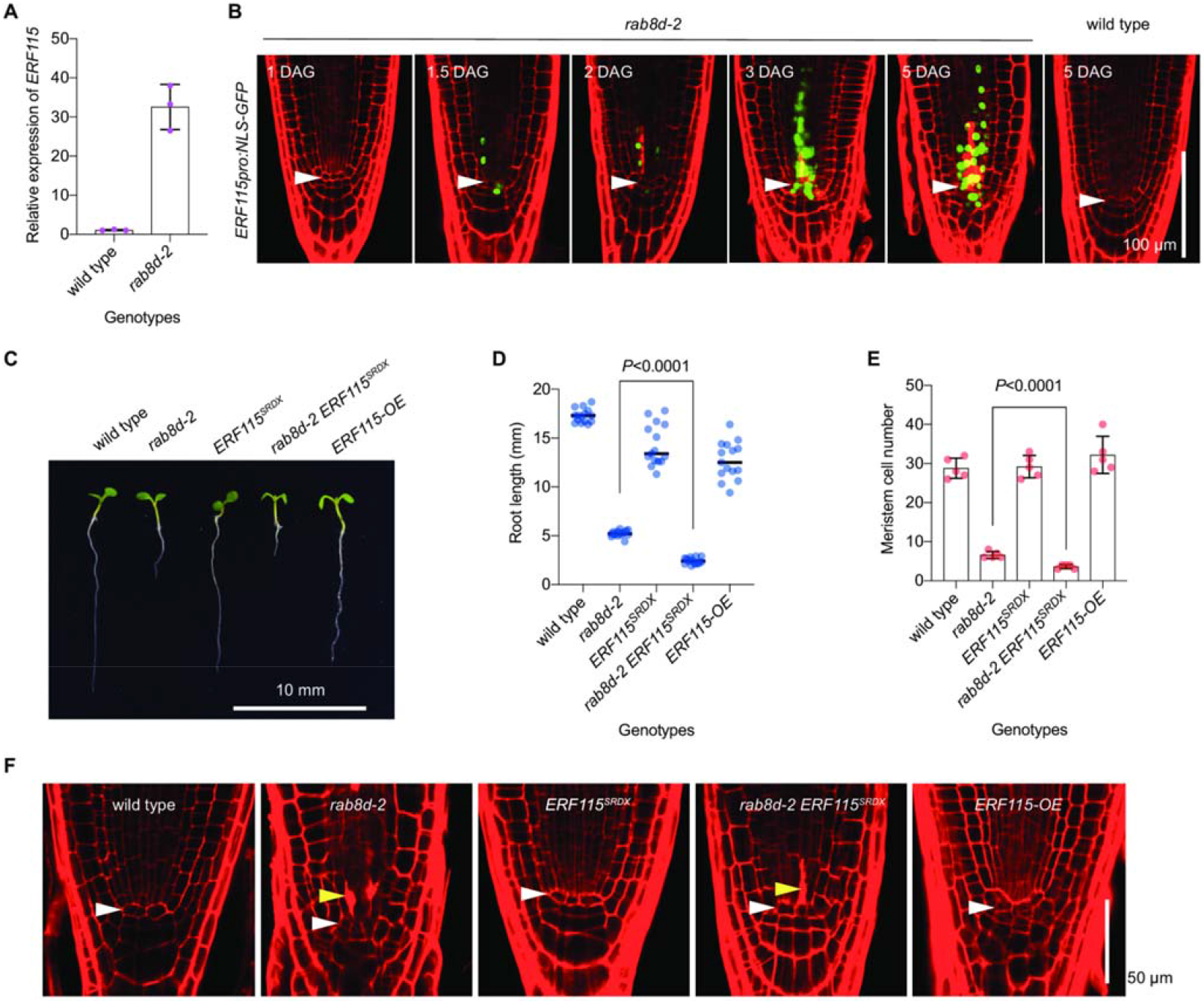
The function loss of *ERF115* inhibits the QC division in *rab8d-2*. (A) Comparison of expression level of *ERF115* between wild type (set to 1) and *rab8d-2*. The data was based on the FPKM values tested by RNA-seq. (B) The expression of *ERF115pro:NLS-GFP* in root tip of *rab8d-2* (from 1 to 5 DAG) and wild type (5 DAG). White arrowheads indicate QC. (C) Phenotypes of 5 DAG seedlings of the indicated genotypes. (D, E) Comparison of root length (D) and meristem cell number (E) among the indicated genotypes. Each spot represents an individual value. For (D), black dashes indicate mean values. For (E), data indicate mean ± SD. Significance analysis was performed using *t* test. (F) SCN phenotypes of 5 DAG seedlings of the indicated genotypes. White arrowheads indicate QC, and yellow ones indicate dead stem cells.

It has been reported that the expression of *ERF115* leads to division of the QC cells (Heyman et al., 2013). Our results also clearly indicated the high expression of *ERF115* in the QC and QC-like cells of *rab8d-2* (Figure 5B). We thus wondered that whether the loss of QC identity in the *rab8d-2* root was associated with the induction of *ERF115*. To address this issue, we crossed *rab8d-2* with a dominant-negative mutant *ERF115^SRDX^*. The results showed that, interestingly, the loss of *ERF115* function further inhibited the root growth of *rab8d-2* (Figure 5C, D). In consistent with this result, the meristem cell number of *rab8d-2 ERF115^SRDX^* was significantly less than that of *rab8d-2* (Figure 5E). More importantly, although the PCD phenotype was observed in SCN, the QC cells appeared to be inactive and were easily identified in *rab8d-2 ERF115^SRDX^* (Figure 5F). In contrast, although the SCN was normal in the overexpression line of *ERF115 (ERF115-OE*), its QC was indistinguishable because of cell division (Figure 5F). These observations suggest that ERF115 has no roles in controlling the longevity of SCN but functions in the activation of QC cells in *rab8d-2*.

### Rab8d physically interacts with GUN1 *in vivo*

To further dig out the molecular mechanism of *rab8d*-dependent programmed death of stem cells, we performed a GFP-tag based co-immunoprecipitation (co-IP) experiment for isolating potential interaction partners of Rab8d from a homozygous line of *Rab8dpro:Rab8d-GFP*. After peptide identification by mass spectrometry (MS), a total of 344 proteins were identified (Figure 6A, Supplemental Dataset 5). As expected, most of these proteins were localized in plastids according to the information from TAIR. In terms of the Eukaryotic Orthologous Groups (KOG) classification, 83 proteins were involved in translation, ribosomal structure and biogenesis, and 55 proteins were involved in posttranslational modification, protein turnover and chaperones (Supplemental Dataset 6), suggesting that Rab8d played a role in maintaining plastid proteostasis. This observation reminded us that another protein, GUN1, also regulates plastid proteostasis through the transient formation of various protein complex with different substrates (Tadini et al., 2016; Marino et al., 2019; Wu et al., 2019). We thus wondered that whether Rab8d and GUN1 shared common partners. Through comparing the proteomic data of GUN1-GFP substrates at the seedling stage from a recent study (Wu et al., 2019) with our data of Rab8d-GFP ones, remarkably, more than one quarter of the identified Rab8d partners (a total of 101) were associated with GUN1 (Figure 6A, Supplemental Dataset 7).

**Figure 6.**
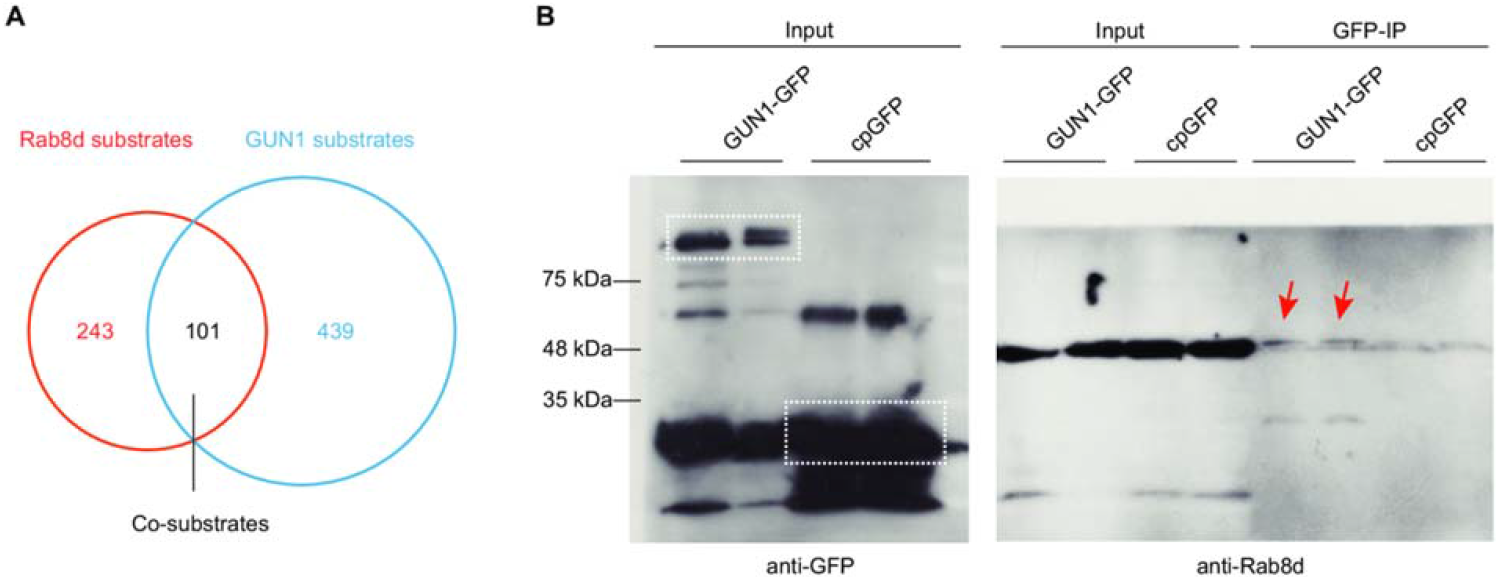
Rab8d physically interacts with GUN1 *in vivo*. (A) Venn diagram indicates the common substrates (co-substrates) between Rab8d and GUN1. The Rab8d substrates were isolated from the *Rab8dpro:Rab8d-GFP* transgenic plants in the *rab8d-2* background. The published data for the GUN1 substrates were employed. (B) Co-IP analysis indicates the interaction between GUN1 and Rab8d *in vivo*. Dotted boxes indicate the bands of pray proteins. Red arrows indicate the bands of Rab8d.

Like Rab8d, the GUN1 protein shows a diffuse distribution pattern in plastids (Wu et al., 2018). The above results implied an association between Rab8d and GUN1 *in vivo*. We did not detect any GUN1 peptides in the Rab8d substrates, which might be resulted from the extremely low protein abundance of GUN1 *in vivo* (Wu et al., 2018). However, based on the published proteomic data (Tadini et al., 2016; Wu et al., 2019), Rab8d presents in the GUN1-dependent substrates. To further confirm the interaction between Rab8d and GUN1, we employed an overexpression line of GUN1-GFP (OE-13) and conducted the GFP-based IP. The result showed that Rab8d could be detected in the direct substrates of GUN1 (Figure 6B). In contrast, the Rab8d level in the cpGFP (a plastid targeted GFP fused with the RbcS transit peptide) substrates was extremely weak (Figure 6B). The results suggest that Rab8d can directly target GUN1 *in vivo*.

### GUN1 determines the longevity of SCN in *rab8d*

Since Rab8d physically interacted with GUN1, we wondered the effect of loss of GUN1 function on the *rab8d* phenotype. To address this issue, we tried to eradicate the GUN1 function in *rab8d-2* by crossing it with *gun1-101* (a widely used T-DNA insertion mutant of *GUN1*). However, no viable homozygous double mutant of *gun1-101 rab8d-2* was isolated. Nevertheless, we found that a small portion of seedlings of the F2 population was etiolated or albino with severely arrested root growth (Figure 7A). These plants could not develop true leaves, suggesting a cotyledon-lethal phenotype (Figure 7A). We further confirmed that these plants were homozygous *gun1-101 rab8d-2* using PCR detection. Notably, the root architecture of double mutant was severely disorganized, the SCN was hardly identified, and many of the root tip cells were enlarged (Figure 7B). The cellular enlargement might be associated with differentiation and endoreduplication. To further confirm this observation, we introduced *gun1-8* (a site-mutation allele of *gun1*) into *rab8d-2*, and a similar phenotype was observed in the *gun1-8 rab8d-2* double mutant (Figure 7B). These results suggest that GUN1 likely is vital for maintaining the root architecture of *rab8d-2*.

**Figure 7.**
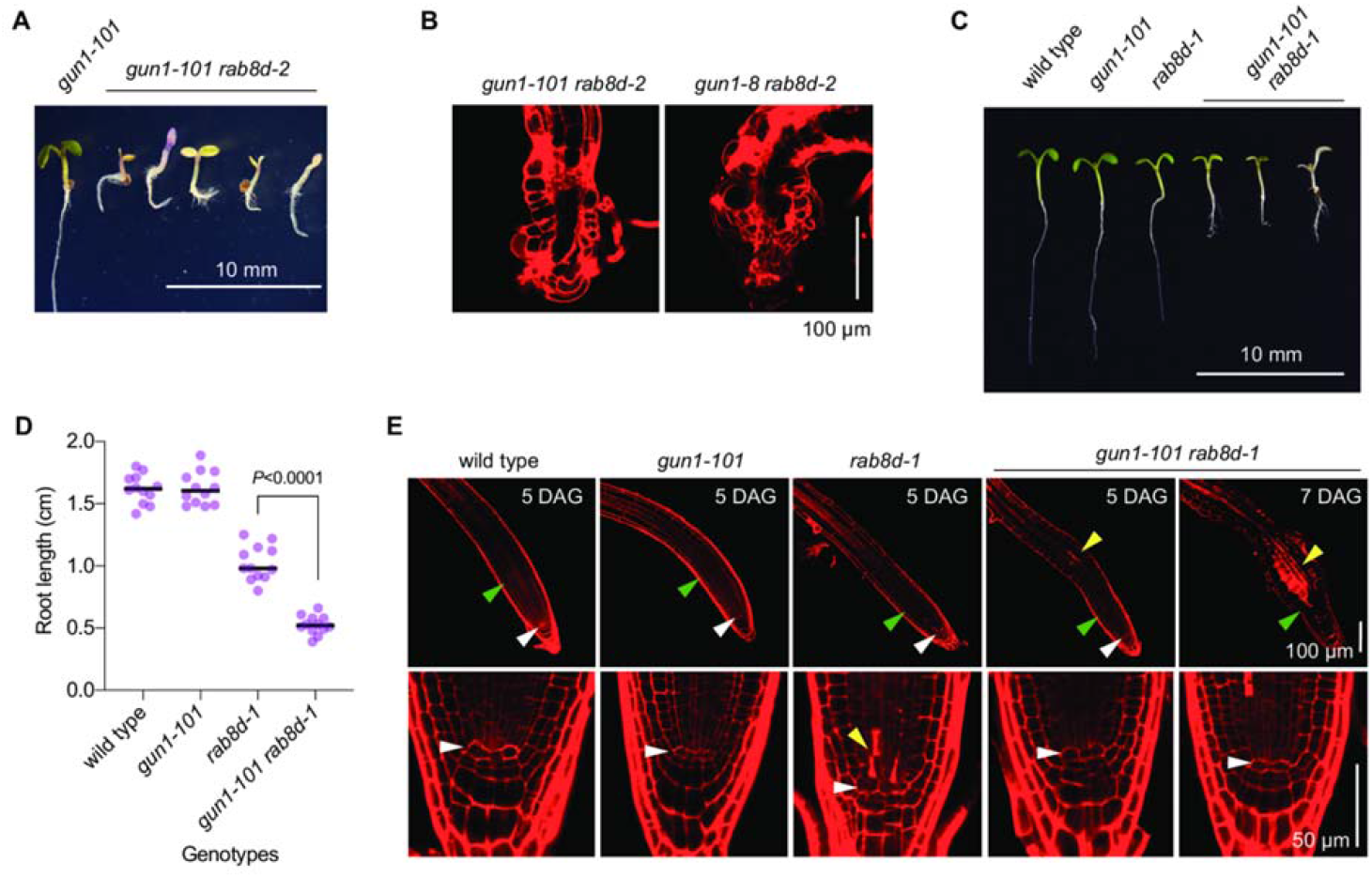
The function loss of GUN1 inhibits PCD in root stem cells of *rab8d-1*. (A) Phenotypes of the *gun1-101 rab8d-2* double mutant at 5 DAG. (B) Phenotypes of root tips of the *gun1-101 rab8d-2* and *gun1-8 rab8d-2* double mutants at 5 DAG. (C) Seedling phenotypes of the indicated genotypes at 5 DAG. (D) Comparison of root length among the indicated genotypes. Each spot represents an individual value, and black dashes indicate mean values. Significance analysis was performed using *t* test. (E) Phenotypes of root tips of the indicated genotypes at the indicated stages. White arrowheads indicate QC, green ones indicate the basal cell of elongation region, and yellow ones indicate dead cells. Note that, at 7 DAG, a large area of cells was dead in the transition/elongation zone.

We assumed that the severe developmental defect of *gun1 rab8d-2* might be partially associated with the very low level (about 8%) of *Rab8d* transcripts in *rab8d-2*. To test this, we crossed *gun1-101* with the *rab8d-1* allele that retained about 30% of *Rab8d* transcripts as mentioned above and analyzed the double mutant phenotypes. The results showed that, indeed, the *gun1-101 rab8d-1* double mutant displayed a mitigated phenotype relative to *gun1-101 rab8d-2* because it could develop true leaves (Figure 7C). Nevertheless, the *gun1-101 rab8d-1* plants were still pale-green or albino and arrested before the reproductive stage (Figure 7C). In addition, the root of *gun1-101 rab8d-1* was significantly shorter than the single mutants (Figure 7D), suggesting that *rab8d-1* was hypersensitive to *gun1-101*, and vice versa. Using PI staining, we found that the loss of *GUN1* function had no clear effect on the root tip architecture, whereas *rab8d-1* displayed phenotypes including the reduced root meristem size, PCD of stele initials, and disorganized QC (Figure 7E). By contrast, at 5 DAG, although the *gun1-101 rab8d-1* double mutant showed a reduced root meristem size phenotype as well, no cell death was observed in the root stele initials, and the QC organization was normal (Figure 7E). These observations suggest that the loss of GUN1 function rescued the abnormal SCN phenotype but had no effect on the meristematic zone of *rab8d-1* at this stage. However, unexpectedly, in the root transition zone of *gun1-101 rab8d-1*, some of the stele cells were filled with PI (indicated by a yellow arrowhead in Figure 7E), suggesting PCD of these cells. At 7 DAG, there was still no cell death in the SCN of *gun1-101 rab8d-1*, but, noticeably, its transition zone was enlarged with death of a large area of cells (Figure 7E). At about 12 DAG, the root architecture of *gun1-101 rab8d-1* was severely impaired, like *gun1-101 rab8d-2* (Supplemental Figure 10). These results suggest that GUN1 likely is required for the death and renewal of root stem cells in the *rab8d* mutants, which might be important for maintaining the root architecture. The observations on *gun1 rab8d-2* also could be explained by this hypothesis.

GUN1 has been well-studied because it is the hub integrating multiple plastid-to-nucleus retrograde signaling pathways (Pesaresi and Kim, 2019; Zhao et al., 2019). Since the mis-regulated expression of a number of nuclear genes by the reduced function of Rab8d, we assumed that the GUN1-dependent plastid signals might be involved in the PCD of root stem cells in the *rab8d* mutants. A recent study has reported that blocking the cytosolic HSP90 function with its specific inhibitor geldanamycin (GDA) significantly mitigated the GUN phenotype of *gun1* (Wu et al., 2019). According to this result, we further tested the phenotypes of *gun1-101 rab8d-1* and *gun1-101 rab8d-2* grown under 50 μM and 100 μM GDA conditions. The results showed that the GDA treatment had no effect on phenotypes of these two lines (Supplemental Figure 11), suggesting that the *rab8d*-dependent signaling was HSP90-independent.

Another recent study reported that GUN1 exerts its role in communication between plastids and nuclei through controlling the tetrapyrrole metabolism (Shimizu et al., 2019). To test whether the tetrapyrrole-related signals were responsible for the PCD of root initials in *rab8d*, we crossed another *gun* mutant, *conditional chlorina* (*cch1-1*, a *gun5* allele), with *rab8d-2*. *GUN5* encodes the H-subunit of Mg-chelatase, which participates in the tetrapyrrole biosynthesis pathway (Adhikari et al., 2011). Unlike *gun1-101 rab8d-2*, the *cch1-1 rab8d-2* double mutant was viable (Supplemental Figure 12). The shoot of *cch1-1* displayed a paler phenotype compared with *rab8d-2*, but its root length was similar to wild type (Supplemental Figure 12A, B). By contrast, the *cch1-1 rab8d-2* double mutant displayed a *cch*-like shoot and a *rab8d*-like root length (Supplemental Figure 12A, B). Importantly, the root tip architecture of *cch1-1 rab8d-2* was also similar to the *rab8d-2* single mutant (Supplemental Figure 12C), which indicated that *rab8d-2* was not sensitive to the loss of GUN5 function. These observations further suggest that the tetrapyrrole-mediated retrograde signaling might not be associated with the PCD phenotype in the root SCN of rab8d.

## DISCUSSION

Plastids, as a kind of semiautonomous organelles in plant cells, are indispensable for normal plant growth and development. Many studies have implied that signals triggered by impaired plastid translation regulate both shoot and root development. For instance, the perturbed function of SCABRA1 (SCA1), a plastid-targeted ribosomal protein, enhances the leaf polarity defects of *asymmetric leaves (as1* and *as2*) mutants (Mateo-Bonmati et al., 2015). Several members of SUPPRESSOR OF VARIEGATIONs (SVRs) have been found to play a role in the leaf margin development (Zheng et al., 2016; Liu et al., 2019). REGULATOR OF FATTY ACID COMPOSITION3 (RFC3), also a plastid ribosomal protein, is required for the stem cell patterning in lateral roots (Nakata et al., 2018). In this study, we discovered an unexpected role for the plastid EF-Tu Rab8d in maintaining the primary root growth and development. Nevertheless, robust inhibition of plastid translation by an exogenous drug lincomycin could not accurately mimic the phenotypes of *rab8d-2* (Supplemental Figure 3). These results suggest that the plastid translation-dependent signaling is sophisticated and multifunctional and may require an accurate signal for regulating root development.

A previous study reported that auxin homeostasis is sensitive to repression of plastid translation in leaves (Zheng et al., 2016). *SVR9* encodes a plastid translation initiation factor, and its function loss leads to disturbed auxin distribution in leaves, which is linked with the abnormal leaf margin development of the *svr9* mutant (Zheng et al., 2016). Rab8d has been identified as an SVR member, SVR11, because its mutation can suppress the variegation phenotype of *yellow variegated (var2*), like other SVRs (Liu et al., 2019). *svr11* enhances the abnormal leaf margin phenotype of *svr9* (Liu et al., 2019). *sca1* also displays a similar phenotype (Mateo-Bonmati et al., 2015). Furthermore, exogenous application of plastid translation inhibitors displays a similar effect on auxin distribution as in the *svr9* mutant (Zheng et al., 2016). These observations suggest an unknown mechanism that plastid translation dependent signals regulate auxin homeostasis in leaves. Our results indicated that the auxin homeostasis was affected in the root tip of *rab8d-2* (Figure 3A), which was tightly associated with the disturbed accumulation of PINs (Supplemental Figure 5). Whether the regulation of auxin homeostasis in roots by *rab8d-2* shares a common mechanism with that in leaves by plastid translation dependent signals needs to be further investigated.

The most interesting phenotype in roots of the *rab8d* mutants is the PCD of stem cells (Figure 1E). This phenotype is frequently observed when DNA is damaged. In both plants and animals, two highly conserved protein kinases ATM and ATM/RAD3-RELATED (ATR) mediate the DNA damage response (DDR), including inhibition of cell cycle progression, promoting DNA repair, PCD, and early endoreduplication (Shiloh, 2006; Cimprich and Cortez, 2008; Hu et al., 2016). In animals, DDR efficiently prevents cancer and protects the germline (Rich et al., 2000). In plants, DDR is also important for the plant viability (Johnson et al., 2018). Mutation in *ATM* or its downstream transcription factor *SOG1* leads to permanent arrest of root growth when plants were suffered severe DNA damage (Garcia et al., 2003; Johnson et al., 2018). Our observations suggest that ATM and SOG1 mediate the inhibition of cell cycle progression in root meristem of *rab8d-2* through inducing the *SMR5* expression, yet, the PCD phenotype in the *rab8d-2* SCN is independent of ATM or SOG1 (Figure 4). We can imagine that if ATM and SOG1 function in PCD in the *rab8d-2* SCN, the double mutants of *atm-2 rab8d-2* and *sog1-1 rab8d-2* should be lethal, like *atm-2* and *sog1-1* under DNA-damaged conditions. Thus, the viability of *atm-2 rab8d-2* and *sog1-1 rab8d-2* is, to some extent, consistent with the independence of the ATM-SOG1 module in the PCD phenotype in the root SCN of *rab8d-2*. These results suggest that the *rab8d*-dependent signaling and the DDR signaling pathways are partially merged. Nevertheless, how ATM perceive the *rab8d*-dependent signal remains elusive. In animal cells, a fraction of ATM targets to mitochondria and exerts its function in modulating mitochondrial homeostasis (Maryanovich et al., 2012; Valentin-Vega et al., 2012). However, the exact subcellular localization of the plant ATM is still mystery until now. A recent study has suggested the mitochondrial localization of Arabidopsis ATM through expressing its N-terminal region fused with GFP (Su et al., 2017), but this observation does not represent the true localization of the full protein. Notably, the MS results of two previous studies from different labs showed that ATM, at least a portion of proteins, targets to the plastid stroma (Kleffmann et al., 2004; Peltier et al., 2004). Thus, the merge of localization between ATM and Rab8d further supports our hypothesis that ATM can perceive the *rab8d*-dependent plastid signal. Yet, the perception might be indirect because ATM is absent from the Rab8d substrates.

The PCD in SCN might be an inducement for the QC division in *rab8d* mutants, because the latter was lagged the former (Figure 1E, F). The expression level of *ERF115* in *rab8d-2* was positively associated with the damage degree of SCN (Figure 5B), which was in agreement with the role of ERF115 in full SCN recovery (Heyman et al., 2016). Consistent with the induction of *ERF115*, the transcriptional level of a target gene of ERF115, *PSK5*, was also increased significantly in *rab8d-2* (based on the RNA-seq data presented in Supplemental Dataset 1). We further revealed that ERF115 was responsible for the QC division in *rab8d-2* (Figure 5F). Because the replenishment of stem cells requires QC division, the loss of ERF115 function might reduce the renewal ratio of stem cells in *rab8d-2*. This hypothesis can explain the much shorter root of *ERF115^SRDX^ rab8d-2* than that of *rab8d-2* (Figure 5C).

The physical interaction between GUN1 and Rab8d suggests that GUN1 can directly monitor the Rab8d homeostasis. It is unexpected that the loss of GUN1 function rescues the PCD phenotype in the *rab8d-1* SCN (Figure 7E) because no studies reveal the role of GUN1 in root development until now. The severely impaired root architecture in *rab8d gun1* double mutants (Figure 7B, E) suggests that the programmed death and renewal of stem cells in *rab8d* mutants is required for maintaining root development. GUN1 is a pentatricopeptide repeat (PPR) protein with a high turnover rate and only can be detected, but still weakly, at the first days after germination (Wu et al., 2018). Under normal growth conditions, the *gun1* mutant displays a slightly delayed plastid differentiation phenotype (Wu et al., 2018). When plastid biogenesis is affected, however, the coordination of gene expression between plastid and nucleus is impaired in *gun1* (Susek et al., 1993). Recent studies have provided two different hypotheses on the mechanism of GUN1-mediated communication between these two organelles. One is dependent on the accumulation of plastid preproteins in cytoplasm resulted from the impaired plastid protein import (Wu et al., 2019), another is associated with changes in tetrapyrrole metabolism (Shimizu et al., 2019). However, none of these two pathways is involved in the PCD phenotype in the *rab8d* SCN (Supplemental Figures 11 and 12).

Together, we raised a model for illustrating the *rab8d*-dependent root phenotype (Figure 8). In the root meristem region, ATM perceives the plastid signal triggered by the reduced function of Rab8d and affects the cell cycle through regulating the *SMR5* expression. Meanwhile, in the root SCN, GUN1 perceives the *rab8d*-dependent signal and triggers PCD through an unknown pathway. The injured SCN induces the expression of *ERF115* for replenishment of stem cells. Thus, our observations imply a crucial role for plastid translation dependent signals in regulating root growth.

**Figure 8.**
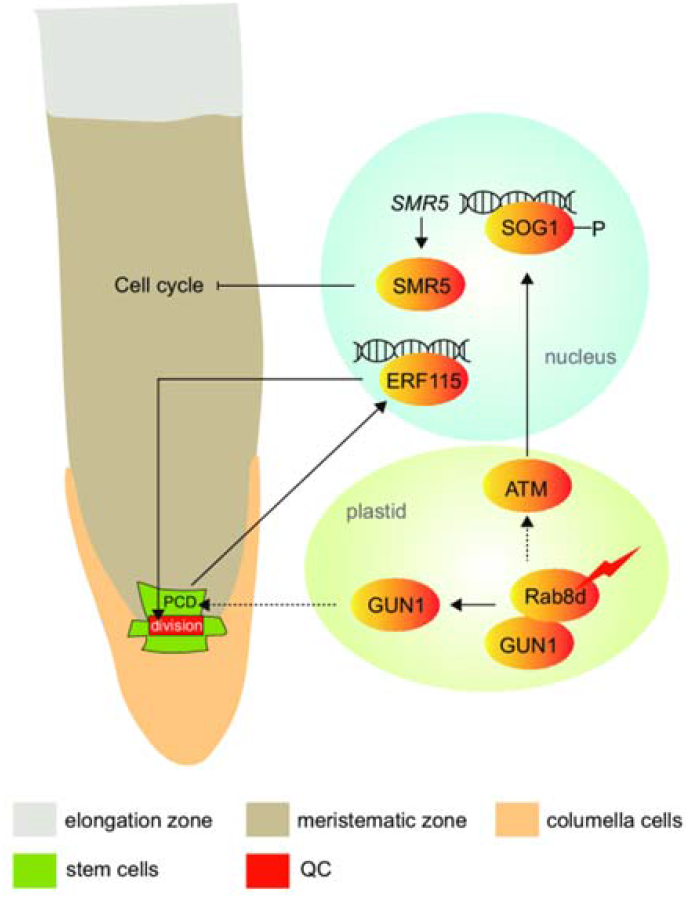
A model for the *rab8d*-dependent signal in regulating root growth. In plastids, when the function of Rab8d is impaired, the derived signal can be perceived by ATM. The activation of ATM signaling phosphorylates a NAC transcription factor, SOG1, which targets to the *SMR5* gene and activates its expression in the root meristem region. The induced SMR5 further inhibits the cell cycle. Meanwhile, GUN1 perceives the impaired Rab8d function and triggers the programmed death of stem cells. The expression of another transcription factor encoded by *ERF115* is highly induced by the injured SCN, which further stimulates the QC division for the replenishment of stem cells. Both GUN1 and ERF115 are vital for the renewal of stem cells. Dotted lines indicate unknown pathways.

## MATERIALS AND METHODS

### Plant materials and growth conditions

In this study, wild type is the *Arabidopsis thaliana* Columbia (Col-0) ecotype except for *gun1-8*, which was generated from *Col6-3* (Zhao et al., 2019). Mutants of *rab8d-1* (sail_659_g09), *rab8d-2* (salk_069644), *rab8d-3* (sail_415_e05), *gun1-101* (sail_33_d01), and *cch1-1* (CS6499) were obtained from Arabidopsis Biological Resource Center. Mutants and transgenic marker lines of *WOX5pro:GFP* (Blilou et al., 2005), *cpGFP* (Wu et al., 2018), *DR5rev:GFP* (Friml et al., 2003), *SHRpro:SHR-GFP* (Nakajima et al., 2001), *SCRpro:SCR-GFP* (Sabatini et al., 2003), *PLT1pro:CFP* (Kutschmar et al., 2009), *PLT2pro:CFP* (Kutschmar et al., 2009), *SMR5pro:GUS* (Yi et al., 2014), *smr5* (Yi et al., 2014), *smr7* (Yi et al., 2014), *atm-2* (Garcia et al., 2003), *sog1-1* (Yoshiyama et al., 2009), *CycB1;1pro:GUS* (Su et al., 2017), *ERF115^SRDX^* (Heyman et al., 2013), *ERF115-OE* (Heyman et al., 2013), and *ERF115pro:NLS-GUS/GFP* (Heyman et al., 2016) have been described previously.

For the construction of *Rab8dpro:Rab8d-GFP* vector, the genomic DNA of *Rab8d* contained the encoding region (TGA free) and its upstream promoter (1510 bp) was cloned into the *pCAMBIA2300* empty vector. Then *GFP* was inserted at the 3’-end of *Rab8dpro:Rab8d*. The primers were listed in Supplemental Table 1. The *Rab8dpro:Rab8d-GFP* construct was introduced into wild type and *rab8d-2* using the *Agrobacterium tumefaciens* (GV3101)-mediated floral dip method. For selection of transgenic lines, 50 μg/mL kanamycin was applied.

Surface-sterilized seeds were plated on solid 1/2 Murashige and Skoog (MS) medium (pH 5.7) containing 1% (w/v) sucrose. After stratification at 4°C for 3 d, the plates were transferred to a phytotron with 22°C and 16-h light/8-h dark conditions as described previously (Li et al., 2019). For the heat stress treatment, seedlings at 2, 3, or 5 DAG grown under normal growth conditions were transferred to an illumination incubator with 37°C and incubated for 4 h or with 30°C and incubated for 3 d according to the experimental demands. For the lincomycin treatment, seeds were plated on the medium described above containing 220 mg/L lincomycin (Sigma Aldrich).

### Microscopy

For observing the root morphology, roots were stained with PI (5 μg/mL) for 5 min and then transferred to the slides for microscopy. The observation methods for embryos and detection fluorescence signals have been described in our previous study (Li et al., 2019). The fluorescence was detected with a CLSM (FV1200, OLYMPUS). For the detection of PI signal, excitation wavelength of 555 nm and emission wavelength of 610 nm were used. For the detection of GFP signal, excitation wavelength of 476 nm and emission wavelength of 510 nm were used. For the detection of chlorophyll signal, excitation wavelength of 633 nm and emission wavelength of 710 nm were used. Fluorescence intensity was further analyzed with the software ImageJ (https://imagej.nih.gov/ij/). For taking the pictures of seedlings, a digital camera (Canon 600D) was employed. For obtaining the photos of GUS staining, a Leica DM4000B microscope was used.

### GUS staining

The method for GUS staining has been described previously (Li et al., 2016). The stained samples were incubated in the chloral hydrate solution (chloral hydrate/glycerol/water = 8:1:2, w/v/v) for 5 min and then transferred to the slides for microscopy.

### EdU detection

Seedlings at 5 DAG were incubated with 10 μM EdU in liquid MS medium for 1 h, then fixed with a 3.7% (w/v) formaldehyde solution in phosphate buffered saline (PBS) for 15 min. Fixed samples were washed with 3% BSA in PBS (w/v), treated with 0.5% Triton X-100 in PBS (v/v) for 20 min, then washed with 3% BSA in PBS (w/v) once more. The EdU detection for samples were according to manufacturer’s instructions of a kit (Click-iT™ EdU Alexa Fluor™ 488 Imaging Kit, C10337).

### Phenotype analyses

Root length was measured from the root tip to the hypocotyl/root joint using ImageJ. Root meristem cell number was estimated by counting the cortical cells from the QC to the basal cell of the elongation region.

The intensity of cells at the S phase or the G2/M transition stage was calculated according to the following methods: the number of cells at the S or G2/M phase is divided by the number of meristem cortical cells, and then the quantitative value multiplies 100%.

### RNA extraction, RNA sequencing, and qPCR

RNA extraction was performed as described previously (Li et al., 2019). qPCR was performed on the 7500 Real-Time PCR system (Applied Biosystems) using a TB Green Premix EX Taq II (Tli RNaseH Plus) Kit (Takara). The calculation method for relative gene expression levels were performed as described previously (Li et al., 2019). Primers were listed in Supplemental Table 1. The high-throughput sequencing was performed on the BGISEQ-500 platform (Beijing Genomic Institution, www.genomics.org.cn). The raw data of sequencing can be obtained from Sequence Read Archive of National Center for Biotechnology Information (accession number: PRJNA603637). The significantly differently expressed genes were isolated based on the fold change > 2 and adjusted *P* value < 0.001. The GO enrichment analysis was performed using the BiNGO plug-in of the Cytoscape software.

### CoIP assay

For performing CoIP analysis on Rab8d, total native proteins were isolated from 5 DAG seedlings of a homozygous transgenic line of *Rab8dpro:Rab8d-GFP* using a Minute Total Protein Extraction Kit for Plant Tissues (SN-009; Invent Biotechnologies) according to the user manual. Seedlings were grounded to fine powder in liquid nitrogen, and the powder was solubilized in the native extraction buffer supplied in the kit with addition of Complete Protease Inhibitor Cocktail (Roche). After incubation on ice for 10 min, the solution was centrifuged at 12,000*g* for 5 min at 4°C. The supernatant was collected and subsequently incubated with anti-GFP mAb Agarose (D153-8; MBL) for 120 min at 4°C (at room temperature for IP assays). After being washed with the extraction buffer for six times, the agarose beads were mixed with 2*loading buffer and boiled for 10 min. Protein samples were isolated by gel electrophoresis and digested with the Trypsin enzyme. The digested peptides were separated with a Shimadzu LC-20AD model nanoliter liquid chromatograph and then passed to an ESI tandem mass spectrometer: TripleTOF 5600 (SCIEX, Framingham, MA, USA). The ion source was Nanospray III source (SCIEX, Framingham, MA, USA), and the emitter was a needle (New Objectives, Woburn, MA, USA) drawn from quartz material. For data acquisition, the mass spectrometer parameters were set as follows: ion source spray voltage 2,300V, nitrogen pressure 30 psi, spray gas 15 and spray interface temperature 150 °C. Protein identification was performed with the Mascot Software V2.3 according to the database of *Arabidopsis thaliana* TAIR10. The raw data of mass spectrometry can be obtained from the PRIDE partner repository of the ProteomeXchange Consortium (accession number: PXD017331).

For performing IP assays on GUN1, the overexpression line of GUN1-GFP (OE-13) was employed. The protein samples of GUN1-GFP substrates were obtained according to the above method, separated with 10% SDS-PAGE, and then detected with anti-GFP (1:1000) for the input and with anti-Rab8d (1:500) for the output.

### Antibodies

The GFP commercial polyclonal antibody was obtained from ABMART (P30010, Shanghai, China). The Rab8d antibody was produced in rabbit with the whole protein as the antigen.

### Accession numbers

Sequence data from this article can be found in the Arabidopsis Genome Initiative database under the following accession numbers: *Rab8d*, AT4G20360; *SMR5*, AT1G07500; *ATM*, AT3G48190; *SOG1*, AT1G25580; *SMR7*, AT3G27630; *ERF115*, AT5G07310; *GUN1*, AT2G31400; *GUN5*, AT5G13630.

## SUPPLEMENTAL DATA

**Supplemental Figure 1.** *rab8d-3* is embryo lethal.

**Supplemental Figure 2.** Comparison of expression of the *Rab8d* gene among the indicated genotypes.

**Supplemental Figure 3.** The SCN phenotype treated with lincomycin.

**Supplemental Figure 4.** Rab8d protein aggregation under heat stress *in vivo*.

**Supplemental Figure 5.** Accumulation of PINs in *rab8d-2*.

**Supplemental Figure 6.** The effect of IAA and BR on root growth of *rab8d-2*.

**Supplemental Figure 7.** Comparison of FPKM values for the indicated genes between wild type and *rab8d-2*.

**Supplemental Figure 8.** Cell cycle detection in root tip of *rab8d-2*.

**Supplemental Figure 9.** Comparison of ROS accumulation between wild type and *rab8d-2*.

**Supplemental Figure 10.** Root architectures of *gun1-101 rab8d-1* and *gun1-101 rab8d-2* at 12 DAG.

**Supplemental Figure 11.** GDA effect on phenotype of *gun1 rab8d*.

**Supplemental Figure 12.** The *gun5* effect on phenotypes of *rab8d-2*.

**Supplemental Table 1.** Primers used in this study.

**Supplemental Dataset 1.** Total genes detected by RNA-seq.

**Supplemental Dataset 2.** Differentially expressed genes detected by RNA-seq.

**Supplemental Dataset 3.** GO enrichment analysis on the upregulated genes in *rab8d-2*.

**Supplemental Dataset 4.** GO enrichment analysis on the downregulated genes in *rab8d-2*.

**Supplemental Dataset 5.** Rab8d substrates detected by CoIP.

**Supplemental Dataset 6.** KOG enrichment list of the Rab8d substrates.

**Supplemental Dataset 7.** The common substrates between Rab8d and GUN1.

## ACKNOWLEDGEMENTS

The authors sincerely thank Zhaojun Ding for sharing *PIN1pro:PIN1-GFP, PIN2pro:PIN2-GFP, PIN3pro:PIN3-GFP, PIN7pro:PIN7-GFP, SCRpro:SCR-GFP, SHRpro:SHR-GFP*, and *DR5rev:GFP* lines, Lieven De Veylder for sharing the *ERF115pro:NLS-GFP, atm, ERF115-OE, SMR5pro:GUS, ERF115^SRDX^, smr5*, and *smr7* lines, Ralph Bock for sharing *cpGFP* and *GUN1-GFP (OE-13*) lines, Kaoru Yoshiyama for sharing *sog1-1*, and Xiaobo Zhao for sharing *gun1-8*. This work was supported by the National Natural Science Foundation of China (Grant No. 31500257), the Agricultural scientific and technological innovation project of Shandong Academy of Agricultural Sciences (CXGC2018E13).

## AUTHOR CONTRIBUTIONS

PL designed the research. PL, JM, and XS performed experiments. PL, JM, XS, CZ, CM, and XW analyzed the data. PL and XW wrote the paper.

